# AUTO-TUNE: SELECTING THE DISTANCE THRESHOLD FOR INFERRING HIV TRANSMISSION CLUSTERS

**DOI:** 10.1101/2024.03.11.584522

**Authors:** Steven Weaver, Vanessa Dávila-Conn, Daniel Ji, Hannah Verdonk, Santiago Ávila-Ríos, Andrew J. Leigh Brown, Joel O. Wertheim, Sergei L. Kosakovsky Pond

## Abstract

Molecular surveillance of viral pathogens and inference of transmission networks from genomic data play an increasingly important role in public health efforts, especially for HIV-1. For many methods, the genetic distance threshold used to connect sequences in the transmission network is a key parameter informing the properties of inferred networks. Using a distance threshold that is too high can result in a network with many spurious links, making it difficult to interpret. Conversely, a distance threshold that is too low can result in a network with too few links, which may not capture key insights into clusters of public health concern. Published research using the HIV-TRACE software package frequently uses the default threshold of 0.015 substitutions/site for HIV pol gene sequences, but in many cases, investigators heuristically select other threshold parameters to better capture the underlying dynamics of the epidemic they are studying.

Here, we present a general heuristic scoring approach for tuning a distance threshold adaptively, which seeks to prevent the formation of giant clusters. We prioritize the ratio of the sizes of the largest and the second largest cluster, maximizing the number of clusters present in the network.

We apply our scoring heuristic to outbreaks with different characteristics, such as regional or temporal variability, and demonstrate the utility of using the scoring mechanism’s suggested distance threshold to identify clusters exhibiting risk factors that would have otherwise been more difficult to identify. For example, while we found that a 0.015 substitutions/site distance threshold is typical for US-like epidemics, recent outbreaks like the CRF07_BC subtype among men who have sex with men (MSM) in China have been found to have a lower optimal threshold of 0.005 to better capture the transition from injected drug use (IDU) to MSM as the primary risk factor. Alternatively, in communities surrounding Lake Victoria in Uganda, where there has been sustained heterosexual transmission for many years, we found that a larger distance threshold is necessary to capture a more risk factor-diverse population with sparse sampling over a longer period of time. Such identification may allow for more informed intervention action by respective public health officials.

## Introduction

The use of genomic data to infer and characterize transmission networks of various pathogens has grown in prominence in the past two decades, with applications to a growing list of pathogens, including viruses such as HIV (Paraskevis et al., 2016), hepatitis C virus (HCV) (Murphy et al., 2019b), or influenza A virus (Jombart et al., 2011), and bacteria such as *M*.*tuberculosis* (Mai et al., 2018) or *A*.*baumanii* (Thoma et al., 2022). Notably, genomic surveillance had a prominent role during the COVID-19 pandemic, including the use of sequencing for the study of transmission clusters (von Rotz et al., 2023; Campigotto et al., 2023). Choosing an appropriate genetic distance threshold is an important part of using a molecular transmission network to track the spread of rapidly evolving pathogens (Liu et al., 2020; Rose et al., 2020). This distance threshold defines the degree of genetic closeness between pathogen sequences, isolated from two individuals, required to suggest them as potential transmission partners in the network. Using a distance threshold that is too large can result in a network with many spurious, making it difficult to interpret and analyze. On the other hand, using a distance threshold that is too small can result in a network with too few links, underestimating connections between individuals and making it difficult to accurately track the spread of the disease (Gore et al., 2022).

To enhance the utility of inferred transmission networks, it is important to carefully consider the appropriate distance threshold, *d*. This threshold may vary depending on the specific disease and the context in which it is spreading. For example, a highly contagious acute respiratory illness (e.g., SARS-CoV-2) may require a smaller *d* than a less contagious chronic illness that is primarily spread through direct contact (e.g., HIV-1). Viruses are more amenable to molecular studies compared to bacteria due to their high genetic divergence and compact genomes. Given the relatively high evolutionary rate of RNA viruses detectable genetic fingerprints can be prioritized for epidemiological studies over short time periods(Paraskevis et al., 2016).

For chronic infections such as HIV, the most appropriate genetic distance threshold should be determined according to the characteristics of the epidemic such as the speed of transmission, and the evolutionary rate of the genomic region analyzed (Liu et al., 2020). Sampling density and possible delays between infection and diagnosis should be considered, since samples close to the time of seroconversion are more likely to cluster than samples from well after infection. Lower thresholds will capture the most closely related sequences, while higher thresholds will capture long-term epidemics and chronically infected individuals (Junqueira et al., 2019). Cluster analysis, i.e., identification and analysis of connected network components, in public health has been used for early identification of increased transmission (Oster et al., 2021, 2018), monitoring response to an HIV outbreak (Tumpney et al., 2020; Sizemore et al., 2020; Tookes et al., 2020), evaluating the effectiveness of interventions (Peters et al., 2016; Wang et al., 2015; Liu et al., 2020) or predicting clusters that are most likely to grow in the near future (Erly et al., 2021; Ragonnet-Cronin et al., 2022). This balance can be achieved through careful analysis and consideration of the specific disease and context.

This study introduces AUTO-TUNE, a method that offers a systematic approach to select genetic distance thresholds for molecular HIV transmission network analysis, based purely on the structure of the collected data. By autonomously optimizing clustering metrics derived from pairwise genetic distances, AUTO-TUNE has the potential to improve the accuracy and reliability of network inference, irrespective of data attributes. The AUTO-TUNE methodology’s independence from supplementary data makes it less sensitive to variations in data collection protocols and enhances its adaptability to various contexts, including potentially other viral diseases.

## Methods

Assume that there are *S* aligned genomic sequessnces (full or partial, e.g. the HIV-1 *pol* gene) for a pathogen of interest, each representing the “consensus” circulating viral diversity at the time of sampling in a single infected individual. We shall infer a putative transmission network comprising *S* nodes, and *E* links (edges), where an edge is drawn between a pair of sequences if the genetic distance between them is at or below a threshold *d*. In such a network, there will be 0 ≤ *C < S* connected components with more than one node (clusters), which are the primary object of inference. This network inference strategy is used by HIV-TRACE (Kosakovsky Pond et al., 2018), where the genetic distance is computed using the Tamura-Nei (TN93) (Tamura and Nei, 1993) model, with a variety of options controlling how to deal with ambiguous nucleotide bases; for HIV-1 such bases are informative since they often represent variants co-circulating in the infected individual at the time of sampling at substantial frequencies(Kosakovsky Pond et al., 2009).

We begin by describing an approach to assign a score to each of the choices of *d* in a plausible/informative range of distances. Note that while such a range is continuous, it is sufficient to only consider distance cutoffs that are in the array of pairwise distances between the sequences, as those are the cut-points where one or more additional edges will be added to the network as *d* is increased.

### Scoring Heuristic Procedure

The network threshold selection procedure proceeds as follows (we provide an example in the Results section as well).

1. For each candidate threshold *d*_L_, in increasing order, ranging from the smallest genetic distance in the dataset, up to either the largest distance or a predetermined maximal threshold, we compute two network statistics: *R*_12_, the ratio of the size of the largest cluster to the size of the second largest cluster, and *C*, the number of clusters in the network at this threshold. A cluster is defined as a connected component in the network with at least two nodes.
2. A priority score is assigned to each *d*_L_. This score measures two properties of the threshold: Does *R*_12_ jump at *d*_L_? How far is the number of clusters *C* at *d*_L_ from the maximal number of clusters computed over all threshold values? Let there be *N* overall *d*_L_ candidate values, and assume we are examining the ith candidate, 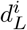 with *W < i* ≤ *N* − *W* (*W* is a positive integer defined below).
  a. The *R*_12_ jump is computed by looking at the normalized ratio of the mean *R*_12_ values computed over the leading window 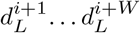 and the trailing window 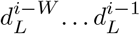. The width of the window, *W*, is defined as 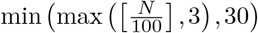. The distribution of ratios is converted to *Z* scores, and normalized relative to the largest positive *Z* score across all candidate distances, yielding the jumeq]p component of the score.
  b. The number of clusters, *C*_i_ at threshold 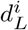 is first normalized to [0, 1] through 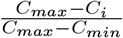 and next gated via a Gompertz function transform 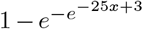. This function provides an *ad hoc* means for penalizing having too few clusters relative to the maximum over all ranges. For example, a threshold that yields 95% of the maximal number of clusters receives a score of 0.996, a threshold that yields 85% - a score of 0.376, and a threshold that yields 60% - a score of 0.0009.
  c. The priority score for 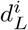 is the sum of the two components defined in (a) and (b), and ranges from 0 to 2.
3. The threshold with the highest priority score will be selected as the suggested automatic distance threshold, if the score is high enough (1.9 or more), and either of the two conditions hold.
  a. No other thresholds have priority scores of 1.9 or higher
  b. If other thresholds have priority scores of 1.9 or higher, then the range of thresholds represented by these options is small (no more than log *N* times the mean step between successive 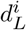).
4. If no single threshold can be selected in step 3, then the one with the highest priority score is suggested, and an inspection of a plot of scores is recommended to ensure that the threshold is sensible.

#### A. Assortativity

Degree-weighted homophily (DWH) is a measure of similarity between nodes in a network based on their attributes (such as demographic characteristics or behaviors) and their degree centrality (i.e., the number of connections they have to other nodes in the network). It is used to quantify the extent to which nodes with similar attributes tend to be connected to each other more frequently than would be expected by chanceGolub and Jackson (2012). DWH is calculated as the ratio of the observed number of connections between nodes with similar attributes to the expected number of connections between such nodes, based on their network degree. For any two subsets *A* and *B* of nodes in a network without singletons (each node has a positive degree), define the weight between *A* and *B* as

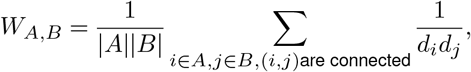

where *d*_i_ is the degree of node *i*, and |*X*| is the cardinality (size) of subset *X*.

Then for any proper (not empty and not the complete network) subset of the network, *G*, e.g. a group of nodes sharing an attribute, e.g., transmission risk factor, define

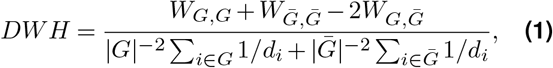

with

- 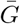: the complement of *G* (all nodes not in *G*)
- *d*_i_ : the degree of node *i*

DWH ranges from −1 to 1. A DWH value of 0 indicates that there is no more homophily than expected by chance (conditioned on network structure), while a value of 1 indicates that there is perfect homophily (*G* consists of connected components disconnected from the rest of the network). A value of −1 is achieved for perfectly disassortative networks (the only links are between *G* and 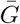).

Homophily metrics have been used in social network analysis and in the study of how different attributes are related to the formation of connections between individuals(Ragonnet-Cronin et al., 2021). To assess whether or not DWH is significantly different from 0 (and from random expectation), we generate the null distribution of DWH obtained by randomly reshuffling node attributes used to define group *G* and recomputing DWH for each such replicate.

#### B. Implementation

The software implementation involves a step-by-step process that utilizes the HIV-TRACE suite of packages. It starts with calculating pairwise distances with the `tn93` tool and a supplied multiple sequence alignment. Thus generated pairwise distances are supplied to the `hivnetworkcsv` script while providing the `-A` keyword argument. A brief outline of the software’s implementation is as follows

1. Calculate pairwise distances: The user first calculates the pairwise distances using the `tn93` fast pairwise distance calculator, providing the maximum threshold value to consider (0.03 in this case, which may be revised upwards for sufficiently divergent sequences, as this provides an upper bound of thresholds to consider) and the input FASTA file. The command for this step is

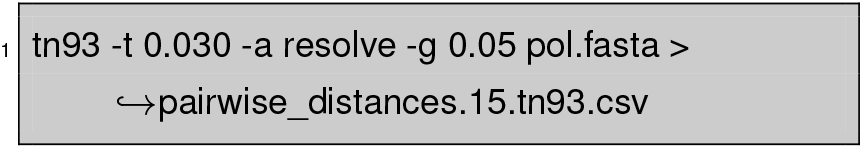 Please note that the threshold should include the maximal range one is intending to test.
2. Compute priority scores for each candidate threshold: The `hivnetworkcsv` script is then executed with the required input file, format, and autotune option to generate a tab-separated output file, as shown below

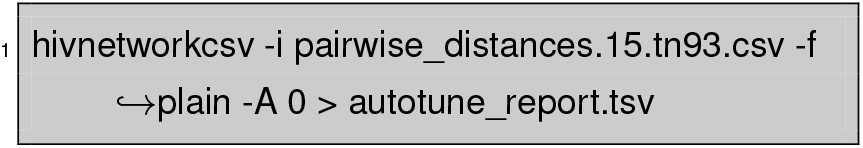
3. Visualize the report: Users can upload the generated `autotune_report.tsv` file to http://autotune.datamonkey.org/analyze for visualization and further analysis of the data. This web-based site extends the Datamonkey platform (Weaver et al., 2018) to provide an interactive environment to explore scores and other metrics across the range of tested outputs.
4. Run HIV-TRACE: Once AUTO-TUNEd threshold(s) are settled upon after review, the user runs the HIV-TRACE command with the appropriate input FASTA file, distance threshold, and other required arguments. The output is saved as a JSON file. An example command is

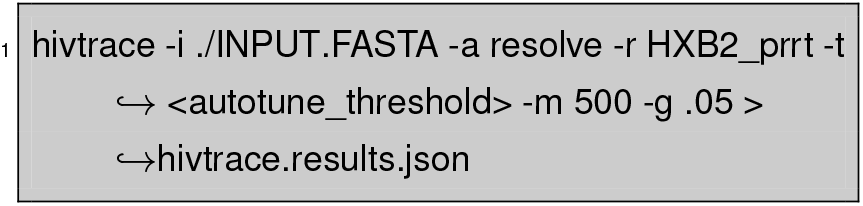

##### B.1. Optional: Compute Assortativity Metrics

5. Annotate results: the `hivnetworkannotate` script is used to annotate the results obtained from the hiv-trace step with attributes. The script takes the JSON results file, node attributes file, schema file, and a resolve flag as input.

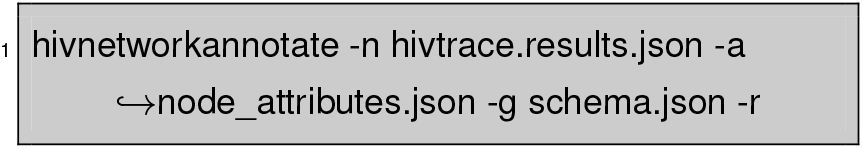 For more information, users can refer to the `hivnetworkannotate` documentation.
6. Analyze the results with DWH: After the results file has been annotated, the user can proceed to the assortativity page, http://autotune.datamonkey.org/assortativity, for further analysis of the output.

The described workflow offers a systematic approach to analyze potential distance thresholds for one’s data with AUTO-TUNE, from calculating pairwise distances to visualizing and annotating results.

#### C. Visualization

Visualizations of AUTO-TUNE results are accessible at http://autotune.datamonkey.org/analyze. These include the priority score plot, and the two contributing statistics: cluster count relative to the maximum and the ratio of two largest cluster sizes. 3 An assortativity tool is available at http://autotune.datamonkey.org/assortativity, and is an analytical tool engineered to facilitate the calculation of Degree Weighted Homophily (DWH) values. It utilizes the DWH NPM package to generate a tabular representation of DWH values corresponding to each value for a selected attribute annotation, providing an exhaustive examination of the interrelationships for the field. The tool also computes the panmictic (null) range, which involves a label permutation test to generate the null distribution of DWH values. This feature establishes a comparative baseline that aids in determining the significance of homophily versus what would be expected by chance.

The visualization code is available on Github (https://github.com/stevenweaver/autotune-app/).

#### D. Comparisons with previously published analyses

First, we set out to compare the thresholds used in numerous published studies with those obtained by AUTO-TUNE. To select the data sets for this analysis, we conducted a scientific literature search to identify studies focused on HIV networks for public health purposes. We then filtered the studies that utilized HIV-TRACE to infer genetic networks and had publicly available sequences. Due to privacy concerns, HIV-1 sequences are frequently not released in the public domain Inzaule et al. (2023). We also attempted to include studies from different countries and regions, enabling us to assess the performance of our method across various epidemic contexts, risk groups, and network sizes in real-data sets that used variable clustering thresholds.

Second, we compared AUTO-TUNE with the most direct published alternative: the `clustuneR` method (Chato et al., 2020). We procured datasets from (Wolf et al., 2017) and (Vrancken et al., 2017) utilizing the identical approach delineated in Chato et al. (2020). These datasets, namely Middle Tennessee, Seattle, and Alberta were processed using the workflow described in Section 2.3. This enabled us to determine an optimal threshold for each dataset using AUTO-TUNE. We further executed the command as detailed in step 4 of Section 2.3, deploying thresholds previously established as optimal by (Chato et al., 2020). Note that `clustuneR` requires and uses temporal information (dates sequences were collected), whereas AUTO-TUNE does not.

Lastly, we evaluated the effect of sampling density on the genetic distance threshold as determined by AUTO-TUNE, we implemented a strategy of random subsampling from the original dataset sourced from (Rhee et al., 2019). This study was selected due to its satisfactory AUTO-TUNE score when utilized in its entirety, as well as its inherent design as a Geographically-Stratified set of 716 *pol* Subtype/CRF (GSPS) reference sequence dataset. The dataset, which comprises 6034 samples gathered between 1989 and 2016, was subjected to random subsampling ten times at proportions of 25%, 50%, and 75% of the original sample size. For each subsample, the optimal threshold and associated scores were determined via AUTO-TUNE.

## Results

### E. Comparisons with published HIV-1 molecular epidemiology studies

We selected several publications citing HIV-TRACE for our analysis, primarily because these studies not only referenced the tool but also made some or all of their sequence data publicly available (Table 1, 2). These studies adopted several different approaches for selecting genetic distance thresholds, including using US CDC guidelines (Yan et al., 2020), picking thresholds based on prior studies (Sivay et al., 2018), and visually inspecting the numbers of clusters and nodes in the networks across candidate distance thresholds (Liu et al., 2020). These thresholds, often qualitatively determined, tended to be round numbers, and were usually determined using *ad hoc* or subjective procedures. Some studies stratified their analyses by viral subtype (major clade), while others did not (or this was not applicable).

**Table 1.**
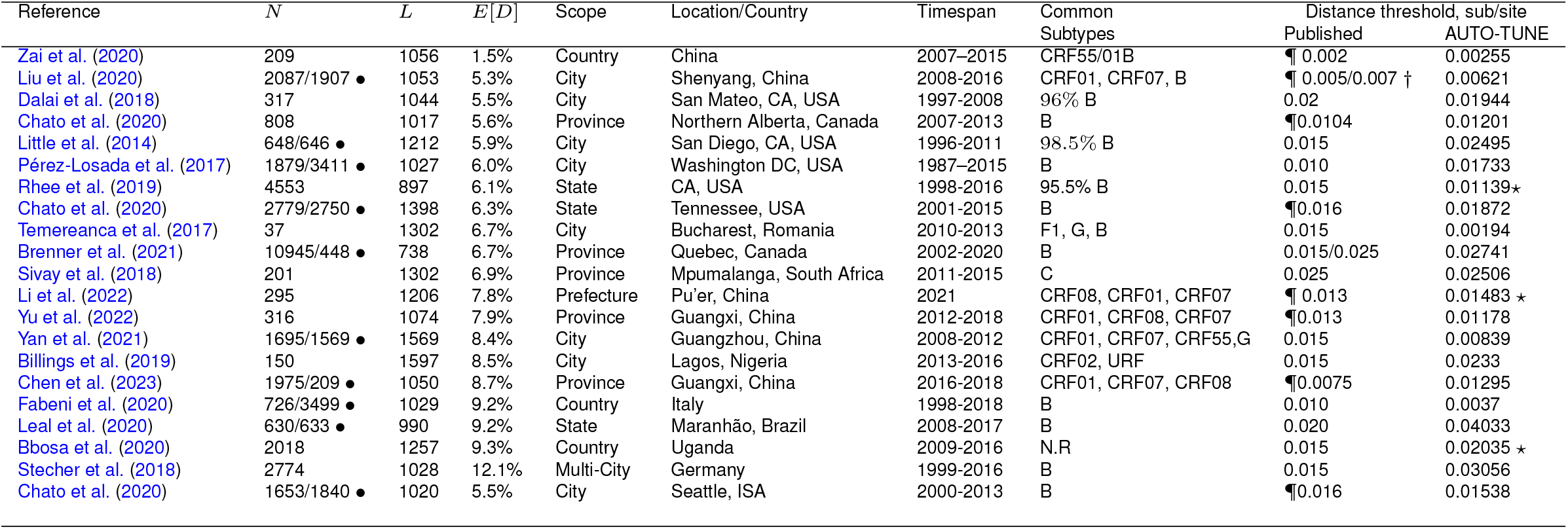
Comparison of AUTO-TUNE and published thresholds from prior studies using partial HIV-1 polymerase gene sequences. *N* : the number of sequences; *L*: length of the multiple sequence alignment, bp; *E*[*D*] mean pairwise TN93 distance; (the studies are sorted on this column, in ascending order) *¶*: the original study performed threshold tuning (varied methods); † : distance thresholds were specific to subtypes; ⋆ : the corresponding AUTO-TUNE score is ≥ 1.9; • : only a subset of the complete dataset was made available (privacy, data use restrictions, incomplete GenBank submissions), the number of sequences analyzed here is shown after the / symbol; N.R: not reported

**Table 2.**
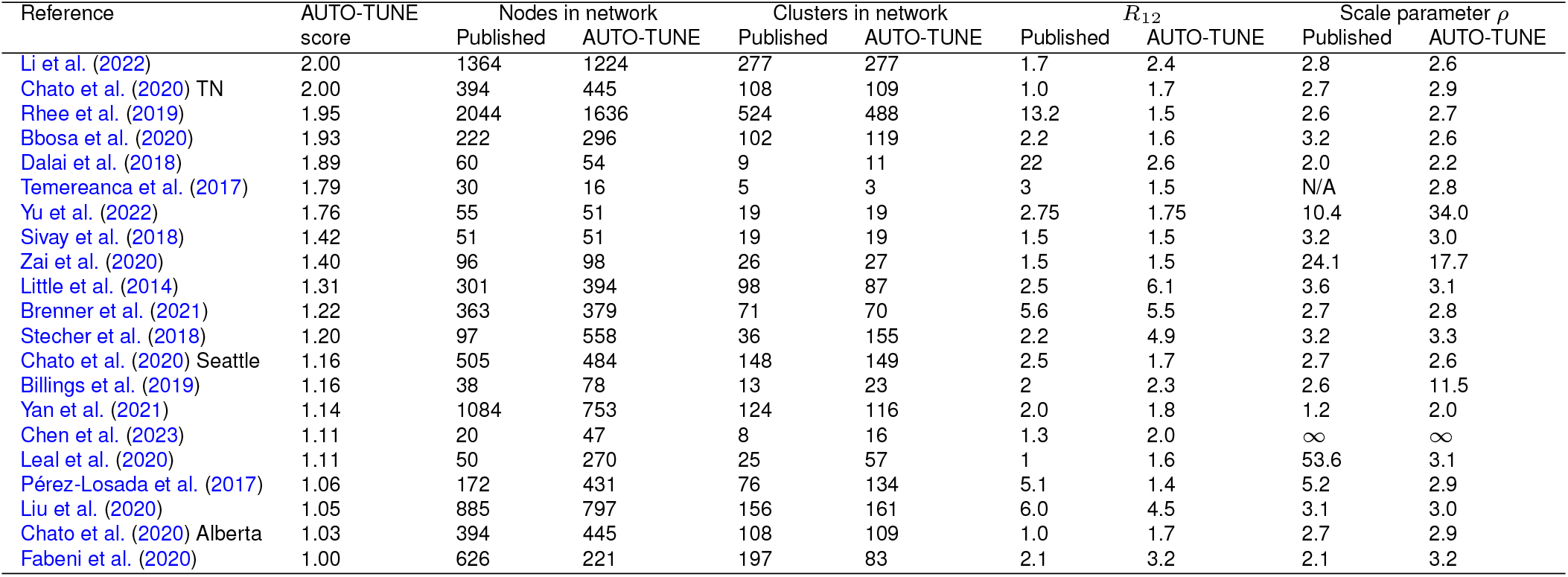
Network properties at the published and AUTO-TUNE thresholds. In cases when the original paper used more than one threshold, we selected the largest for comparison. The datasets are ordered by the AUTO-TUNE priority score from highest to lowest. *ρ* is the fitted characteristic scale-free exponent of the corresponding degree distributions.

| Reference | AUTO-TUNE<br>score | Nodes in network | | Clusters in network | | $R_{12}$ | | Scale parameter $\rho$ | |
| --- | --- | --- | --- | --- | --- | --- | --- | --- | --- |
|  |  | Published | AUTO-TUNE | Published | AUTO-TUNE | Published | AUTO-TUNE | Published | AUTO-TUNE |
| Li et al. (2022) | 2.00 | 1364 | 1224 | 277 | 277 | 1.7 | 2.4 | 2.8 | 2.6 |
| Chato et al. (2020) TN | 2.00 | 394 | 445 | 108 | 109 | 1.0 | 1.7 | 2.7 | 2.9 |
| Rhee et al. (2019) | 1.95 | 2044 | 1636 | 524 | 488 | 13.2 | 1.5 | 2.6 | 2.7 |
| Bbosa et al. (2020) | 1.93 | 222 | 296 | 102 | 119 | 2.2 | 1.6 | 3.2 | 2.6 |
| Dalai et al. (2018) | 1.89 | 60 | 54 | 9 | 11 | 22 | 2.6 | 2.0 | 2.2 |
| Temereanca et al. (2017) | 1.79 | 30 | 16 | 5 | 3 | 3 | 1.5 | N/A | 2.8 |
| Yu et al. (2022) | 1.76 | 55 | 51 | 19 | 19 | 2.75 | 1.75 | 10.4 | 34.0 |
| Sivay et al. (2018) | 1.42 | 51 | 51 | 19 | 19 | 1.5 | 1.5 | 3.2 | 3.0 |
| Zai et al. (2020) | 1.40 | 96 | 98 | 26 | 27 | 1.5 | 1.5 | 24.1 | 17.7 |
| Little et al. (2014) | 1.31 | 301 | 394 | 98 | 87 | 2.5 | 6.1 | 3.6 | 3.1 |
| Brenner et al. (2021) | 1.22 | 363 | 379 | 71 | 70 | 5.6 | 5.5 | 2.7 | 2.8 |
| Stecher et al. (2018) | 1.20 | 97 | 558 | 36 | 155 | 2.2 | 4.9 | 3.2 | 3.3 |
| Chato et al. (2020) Seattle | 1.16 | 505 | 484 | 148 | 149 | 2.5 | 1.7 | 2.7 | 2.6 |
| Billings et al. (2019) | 1.16 | 38 | 78 | 13 | 23 | 2 | 2.3 | 2.6 | 11.5 |
| Yan et al. (2021) | 1.14 | 1084 | 753 | 124 | 116 | 2.0 | 1.8 | 1.2 | 2.0 |
| Chen et al. (2023) | 1.11 | 20 | 47 | 8 | 16 | 1.3 | 2.0 | ∞ | ∞ |
| Leal et al. (2020) | 1.11 | 50 | 270 | 25 | 57 | 1 | 1.6 | 53.6 | 3.1 |
| Pérez-Losada et al. (2017) | 1.06 | 172 | 431 | 76 | 134 | 5.1 | 1.4 | 5.2 | 2.9 |
| Liu et al. (2020) | 1.05 | 885 | 797 | 156 | 161 | 6.0 | 4.5 | 3.1 | 3.0 |
| Chato et al. (2020) Alberta | 1.03 | 394 | 445 | 108 | 109 | 1.0 | 1.7 | 2.7 | 2.9 |
| Fabeni et al. (2020) | 1.00 | 626 | 221 | 197 | 83 | 2.1 | 3.2 | 2.1 | 3.2 |

A direct comparison with published networks is not feasible because only the underlying sequence data (and often only some of the sequences) are made available, not the networks themselves. To facilitate comparison here, we used distance thresholds and all available sequences from primary publications to infer transmission networks anew (the scripts for doing so and the corresponding settings are available in github.com/veg/auto-tune) and compare them with the networks obtained using the highest scoring AUTO-TUNE threshold.

With a few exceptions (e.g Dalai et al. (2018); Sivay et al. (2018)), both the distance thresholds and the inferred networks were quite different, in terms of the numbers of connected nodes, clusters, degree distributions, and even hyper-parameters, such as the characteristic exponent of the scale free degree distribution, *ρ*. This is true even for the studies where the published threshold was tuned (typically to maximize the number of clusters). AUTO-TUNE thresholds were larger than the published values in 13*/*21 datasets, and smaller in 8*/*21 datasets.

### E.1. Examples of how changing thresholds affects inferred networks

#### Cluster size reduction

The 0.02 subs/site (substitutions/site) threshold used by Dalai et al. (2018) yielded one large cluster composed of two loosely connected components (one PWID/HSX, one MSM, see Figure 2 in that paper). A minute change to the threshold by AUTO-TUNE to 0.0194% splits one large cluster into three (some nodes also became disconnected), separating the two major risk groups; this is because the “bridging” connections were between these two thresholds (see Fig 4 panel A). This minor change also reduced *R*_12_ from 21 to 2.6.

**Figure 1.**
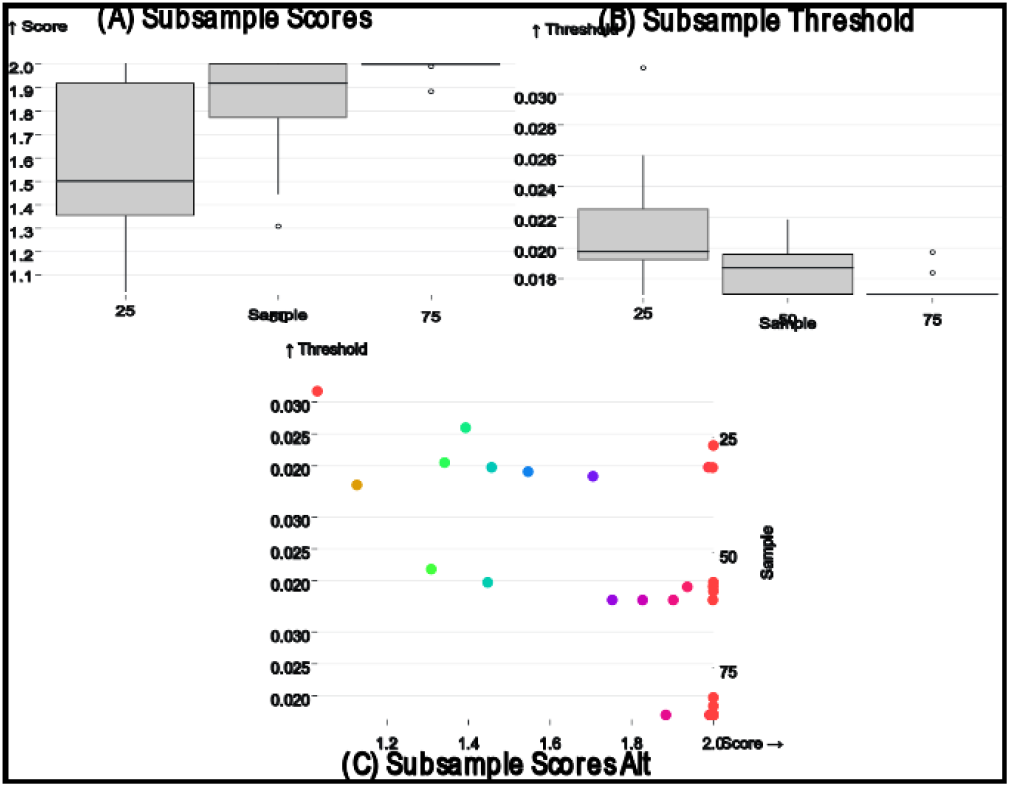
(A) Box plot representing the AUTO-TUNE scores across ten random samples at 25%, 50%, and 75% of the (Rhee et al., 2019) dataset, showing a trend of increasing confidence in score estimates with denser sampling. (B) Box plot of the selected distance thresholds across the same random samples at 25%, 50%, and 75% proportions, demonstrating improved consistency in threshold selection with increased sample size. (C) Scatterplot of the chosen thresholds (Y-axis) against their corresponding AUTO-TUNE scores (X-axis) for the three subsample proportions.

**Figure 2.**
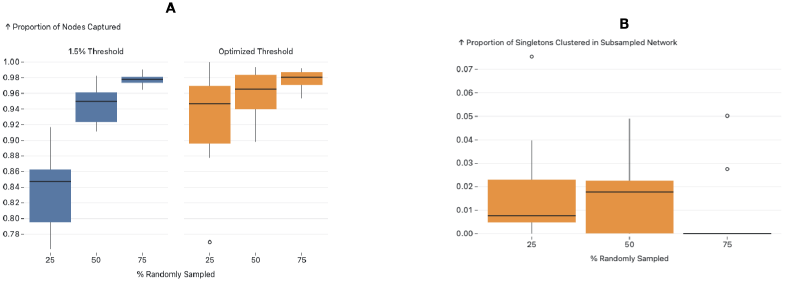
Figure A and B present the effects of subsampling on network structure using different thresholds. Figure A illustrates the proportion of nodes subsampled that remained clustered in both the original and the subsampled networks, with an observable increase in nodes captured as the threshold transitions from 1.5

**Figure 3.**
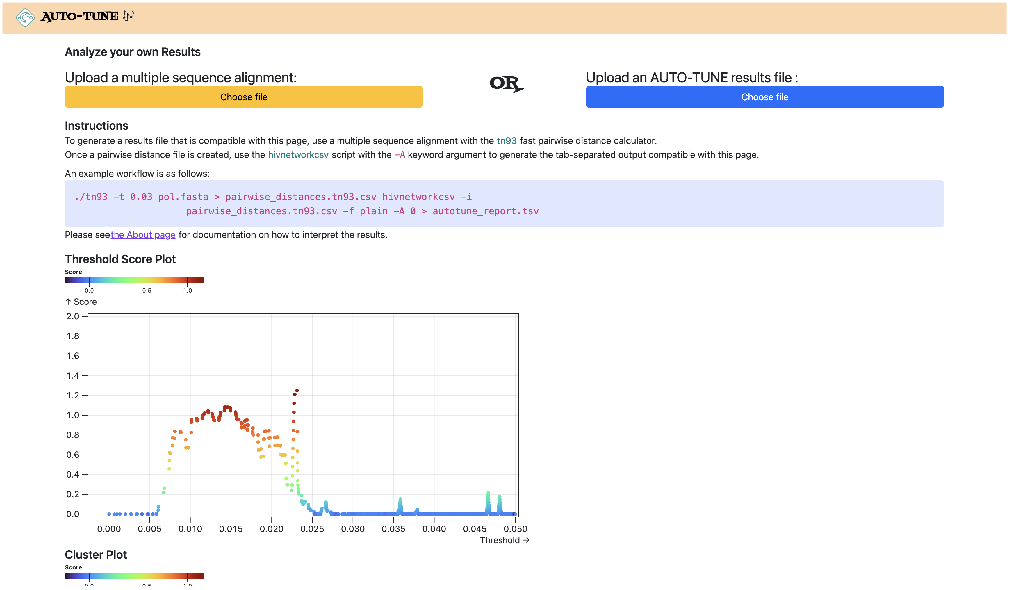
The user interface of the AUTO-TUNE web application (http://autotune.datamonkey.org/analyze). The platform provides a multi-faceted view of AUTO-TUNE’s analysis, including a score plot that visualizes trends across different genetic distance thresholds. It also displays graphs of the number of clusters and the R1/R2 ratio—both key metrics in AUTO-TUNE’s heuristic scoring system. These interactive visualizations aid researchers in making nuanced decisions for threshold selection, especially when multiple thresholds yield similar scores.

**Figure 4.**
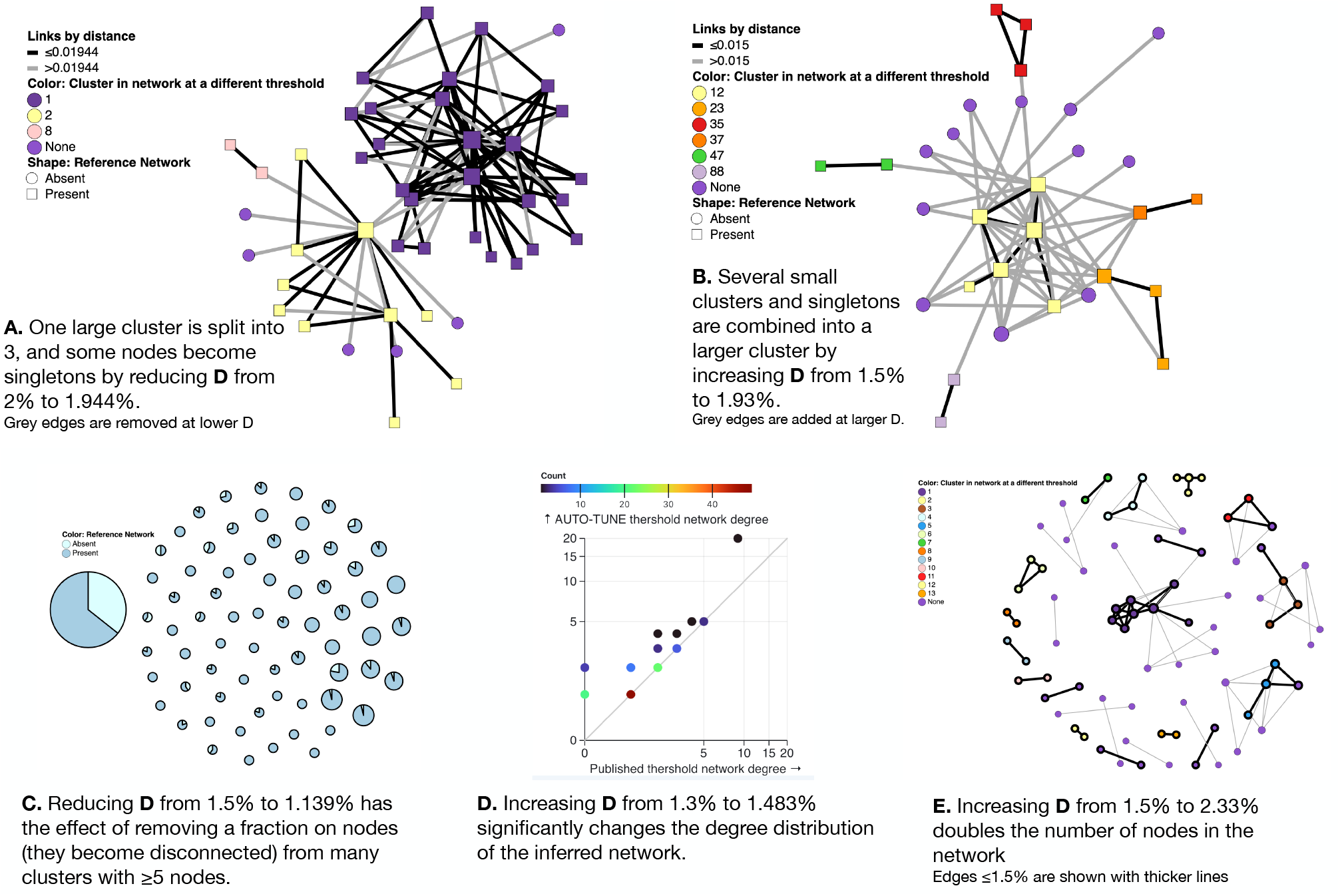
Examples of AUTO-TUNE scores profiles. (A). Lowering the genetic distance threshold removes some of the edges from the network (shown in grey) and disconnects a large cluster into color-coded smaller clusters; here “None” means that the node is not connected to anything at the lower threshold. (B). Raising the genetic distance threshold adds edges to the network (shown in grey) and connectes previously separte clusters into a larger component. (C). Each circle is a cluster in the larger threshold network, and with a proportion of nodes removed when the threshold is lowered. (D). Changes to the node degree distribution (colors represent the counts of nodes with the same degree). (E). A significant enlargement of a small network at a higher threshold, with grey edges only present at the larger threshold.

#### Cluster size increase

Increasing the 0.015% subs/site threshold on data from Little et al. (2014) combined several small clusters (and singletons) into a single larger cluster, while preserving the overall size and properties of the network (see Fig 4 panel B). This change also reduced *R*_12_ from 2.5 to 1.5.

#### Thinning out the network

Reducing the 0.015% sub-s/site threshold on data from Rhee et al. (2019) dramatically reduced the size of the largest cluster, and thinned out most clusters with five or more nodes (see Fig 4 panel C).

#### Materially changing the degree distribution of the network

For the sequences from Li et al. (2022), AUTO-TUNE suggests *D* = 1.483% with robust (1.76) confidence, whereas the original *D* = 0.013 subs/site was selected based on maximizing the number of clusters (and likely rounding to the nearest decimal). While the total number of the clusters only increases by 1, the number of nodes connected in the network grows from 95 to 119, and the scale free exponent of the distribution is dramatically affected. The latter is informed by the degree distribution of the network, and Fig 4 panel D shows, the degree distribution is dramatically affected. Many commonly used network-derived correlates (e.g. degree centrality) can be strongly affected by such changes.

#### Expanding the network

Increasing the .015 subs/site threshold in Billings et al. (2019) to 2.33% more than doubles the number of nodes included (Fig 4 panel E) Networks with high AUTO-TUNE scores are exemplified by the alignment (in the distance space) of the points where the number of clusters is maximized and where the network transitions to having an “unusually” large cluster (see Fig. 5, panel A). In cases of low scores, AUTO-TUNE effectively falls back to maximizing the number of clusters as a function of the distance thresholds, which is a common strategy found in empirical studies (see Fig. 5, panel B).

**Figure 5.**
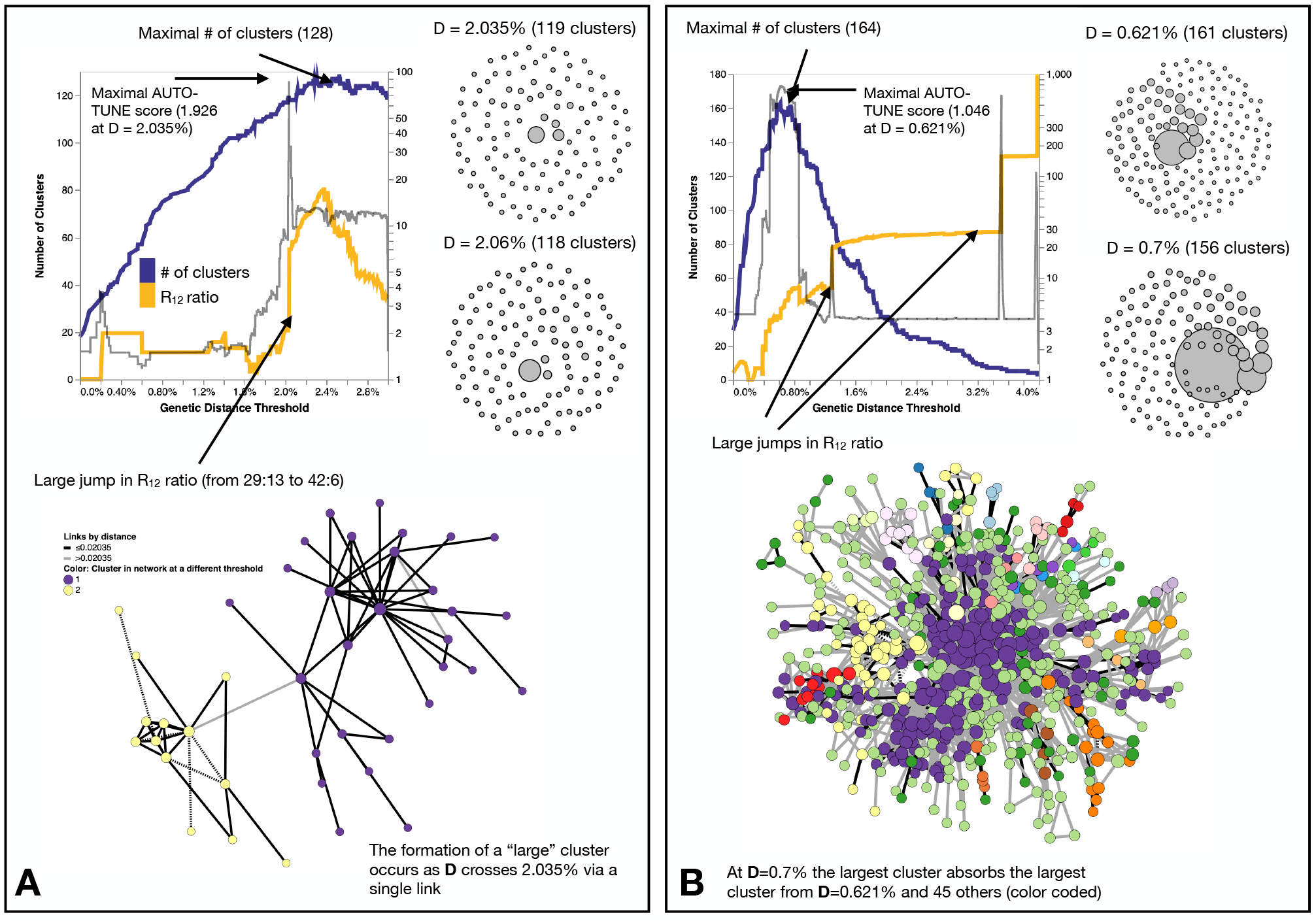
Examples of how changing thresholds affects inferred networks. (A). A high-scoring network Bbosa et al. (2020) has a distance threshold which achieves the number of clusters near the maximum, while also avoiding the formation of a large (weakly connected) cluster. (B). A low-scoring network Liu et al. (2020) has a misalignment between the distance for which the maximum number of clusters is found, and where the big jumps in the cluster size ratio occur. Here, AUTO-TUNE effectively optimizes the number of clusters while preventing excessive growth of the largest cluster.

As expected, AUTO-TUNE inferred smaller thresholds for younger (e.g., studies based in China) epidemics. While AUTO-TUNE will always return a score, in the majority of cases there is no clear “winner” threshold, with priority scores exceeding 1.5 in only 6/18 cases (Table 2). One interpretation for such lack of clarity is that the underlying network has several different (e.g. spatial, temporal, or subtype-specific) thresholds which cannot be well-represented by any single value. For instance, when analyzing the data from Yan et al. (2021), AUTO-TUNE returned a low score of 1.14 for *D* = 0.839%. However, when we split the data into major constituent subtypes and ran AUTO-TUNE on each one separately, starkly discrepant thresholds were found for different subtypes: *D* = 0.0102 subs/site (score = 1.59) for CRF01, *D* = 0.00193 subs/site (score = 2) for CRF05, *D* = 0.02615 subs/site (score = 1.65) for B, and *D* = 0.0111 subs/site (score = 1.04) for CRF07. Although many networks from the literature tend to be dominated by sequences from the same subtype, in more heterogeneous settings it seems prudent to partition the data by subtype (corresponding to major phylogenetic clades), and perform network analyses within subtypes.

### F. Comparisons with published non-HIV molecular epidemiology studies

While HIV-1 epidemiology is the predominant niche for distance-based molecular transmission analyses, other rapidly evolving viruses, especially HCV, have also been analyzed with these approaches(Bartlett et al., 2017). Unlike HIV-1, there is considerably less work on how to choose an appropriate distance threshold, further complicated by the use of different genes to build networks (see Chan et al. (2020) for a comprehensive summary). Two commonly seen methods exist: use some measure of intra-host variation (obtained by deep sequencing) as a lower bound for the threshold, or tune *D* to obtain some desired network property, e.g., the maximal number of clusters. Like with HIV-1, we searched the literature for relevant studies, and selected several with publicly available sequence data.

Most of the datasets are much smaller and less systematically sampled than those for HIV-1, and often combine highly divergent subtypes in the same collection, making a joint analysis challenging. As with HIV-1, AUTO-TUNE returns a wide range of scores and *D* thresholds. For example, effectively maximizing the number of clusters on rhinovirus sequences from Ng et al. (2022) yields a *D* estimate very similar to that obtained by the authors from intra-host variability – information not available to AUTO-TUNE. Table 3

**Table 3.**
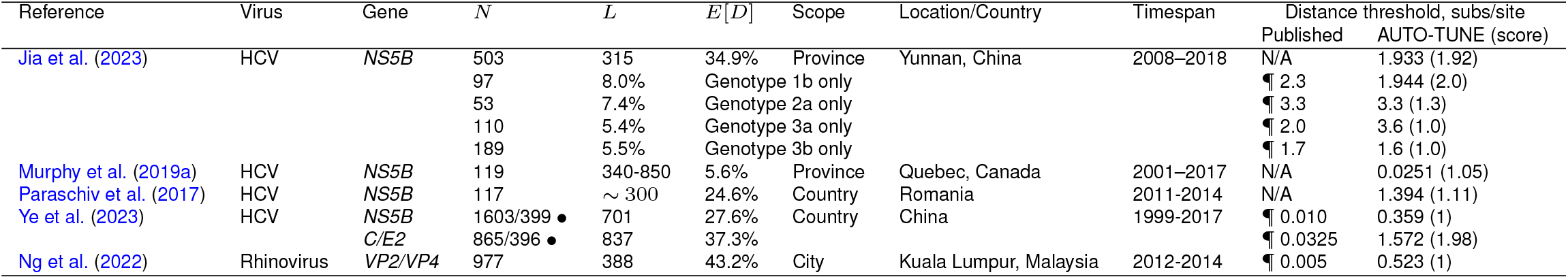
Comparison of AUTO-TUNE and published thresholds from prior studies using sequences from viruses other than HIV-1. “N/A”: no distance-based clustering analyses were done. Other notation is the same as in Table 1

### G. Large-scale HIV-1 database analyses

#### G.1. Markedly different thresholds for different subtypes

Following the spirit of the analysis performed by Wertheim et al. (2014), we downloaded partial *pol* sequences (between HXB-2 coordinates 2253 and 3200, one sequence per patient) from the Los Alamos HIV-1 Database, split them by annotated subtype and applied AUTO-TUNE to individual subtypes with 1000 or more sequences.

Some (but not all) HIV-1 subtypes often act as strong correlates of regional and temporal distributions of sequences, and are expected to represent epidemics with different sampling rates and transmission dynamics. These differences are reflected in a wide range of mean pairwise distances and inferred AUTO-TUNE thresholds shown in Table 4. For example, the relatively young subtype A6, which is the most common subtype in the countries of the former Soviet Union Abidi et al. (2021), has a low mean pairwise distance (0.046) and a low AUTO-TUNE threshold (0.0056). In contrast A1D recombinant sequences have high distance and threshold values (0.089 and 0.0323, respectively), because sequences of this “subtype” represent broadly circulating strains with complex backgrounds, and extensive histories of recombination Foster et al. (2014); Yebra et al. (2015).

**Table 4.** An application of AUTO-TUNE to subtype stratified HIV-1 pol sequences from the LANL database. Fraction clustered is the proportion of all sequences that are connected to at least one other sequence. Subtypes are sorted by the inferred threshold, lowest first. Other notation is the same as in Table 1

| Subtype | $N$ | $E[D]$ | AUTO-TUNE | | Fraction Clustered, % | Mean Degree | $\rho$ |
| --- | --- | --- | --- | --- | --- | --- | --- |
|  |  |  | Threshold, % | Score |  |  |  |
| CRF55 | 2237 | 2.6 | 0.187 | 1.20 | 29.6 | 1.41 | 2.41 |
| CRF07 | 11682 | 3.3 | 0.26 | 1.00 | 26.9 | 3.42 | 1.87 |
| CRF63 | 1649 | 3.6 | 0.505 | 1.01 | 22.1 | 4.85 | 1.8 |
| 01B | 2237 | 7.8 | 0.518 | 1.08 | 22.2 | 0.97 | 5.05 |
| A6 | 11991 | 4.6 | 0.558 | 1.09 | 18.6 | 5.55 | 1.6 |
| CRF08 | 2538 | 3.9 | 0.82 | 1.95 | 25.6 | 1.95 | 2.12 |
| CRF01 | 25689 | 5.1 | 0.875 | 1.73 | 47.0 | 1.94 | 5.54 |
| B | 106261 | 6.4 | 1.084 | 2.00 | 46.4 | 4.77 | 1.95 |
| D | 3561 | 6.6 | 1.133 | 1.12 | 20.8 | 3.65 | 0.79 |
| C | 30714 | 6.7 | 1.438 | 2.00 | 19.3 | 1.26 | 2.22 |
| A1 | 7154 | 7.0 | 1.89 | 2.00 | 17.0 | 5 | 1.7 |
| CRF02 | 7821 | 6.3 | 1.97 | 1.01 | 34.3 | 10.44 | 1.53 |
| BF1 | 4825 | 8.1 | 2.046 | 1.03 | 25.1 | 2.27 | 1.95 |
| G | 2162 | 7.3 | 2.407 | 1.03 | 49.0 | 9.1 | 1.66 |
| F1 | 3986 | 7.6 | 2.941 | 1.34 | 50.4 | 15.03 | 1.40 |
| A1D | 1284 | 8.9 | 3.23 | 1.70 | 27.5 | 1 | 4.3 |
| BC | 2724 | 8.0 | 3.54 | 1.00 | 81.4 | 71.2 | 1.32 |
| Wertheim et al. (2014) | 84527 | . | 1.0 | N/A | 15.6 | 3.84 | 1.74 |

There was extensive variability among subtypes in all high-level network statistics, including the mean degree, fractions of nodes that were in the network, and the characteristic exponent *ρ*, where *ρ* is inferred from by fitting the degree distribution to various network formation models, and with Prob(degree = *k*) ∼ 1*/k*^ρ^ for large *k*.

For A1, B, C, and CRF08 networks there’s very strong support for a single AUTO-TUNE threshold (score *>* 1.9), while for many other subtypes there is extreme ambiguity in which threshold to choose (score *<* 1.1). We suggest that networks where AUTO-TUNE fails to find a single threshold may comprise heterogeneous data which require multiple thresholds to resolve.

#### G.2. Congruence of networks inferred from different genes

Very few published studies of HIV-1 transmission networks use genes other than *pol*, and nearly all of the extrinsically motivated thresholds have been derived for this gene, the utility of other genes and the appropriate *D* values for them are unclear. Because of different rates of evolution in HIV-1 genes and, possibly, subtypes Penn et al. (2008), one would expect *D* to be different for different subtypes and genes. As a simple exercise, we downloaded full-length HIV-1 genomes from the LANL database, stratified them by subtype, and conducted AUTO-TUNE inference using four genomic segments: protease+reverse transcriptase, integrase, matrix (gag), and the less variable gp41 segment of the envelope gene.

Only three subtypes had ≥ 500 full-length sequences in the LANL HIV database (Table 5): B, C, and CRF01. As expected, the inferred thresholds differed by gene and subtype, with lower thresholds inferred for more slowly evolving segments (PR+RT and INT), and similar numbers of clusters found in the resulting subtype-level networks. For all three subtypes, the level of agreement between the four networks on whether or not nodes were clustered or not (present / absent from the network), measured by Krippendorff’s *α* Hayes and Krippendorff (2007), were substantially higher than expected by chance (*α* = 0). All four networks also had between a quarter and a half of all the clusters in perfect agreement.

**Table 5.** Distance thresholds and key network properties using four different HIV-1 genomic regions, stratified by subtype (minumum 500 sequences)

| Subtype | $N$ | AUTO-TUNE $D, \text{subs/site}$ | | | | Number of clusters | | | | Full agreement clusters | Krippendorff $\alpha$ |
| --- | --- | --- | --- | --- | --- | --- | --- | --- | --- | --- | --- |
|  |  | pr+rt | integrase | gag | gp41 | pr+rt | integrase | gag | gp41 |  |  |
| B | 1843 | 0.02081 | 2.0005 | 3.137 | 5.095 | 115 | 128 | 119 | 144 | 64 | 0.723 |
| C | 877 | 0.03266 | 0.02 | 4.754 | 5.325 | 44 | 35 | 47 | 46 | 21 | 0.588 |
| CRF01/AE | 624 | 0.01635 | 0.818 | 2.285 | 2.037 | 40 | 30 | 40 | 41 | 12 | 0.610 |

### H. Evaluating Inferred Networks using Homophily

Non-random mixing and attribute-based homophily are intrinsic characteristics of human contact networks and can be expected to be present in transmission networks, particularly in the context of HIV transmission. People frequently engage in relationships with those who share similar attributes or behaviors, such as risk factors (e.g., PWID, MSM). Recent evidence suggests that race/ethnicity is also a strong predictor of homophily in HIV networks(Ragonnet-Cronin et al., 2021). The effect of these nonrandom mixing patterns on the genetic diversity of HIV-1 has not only been extensively explored via modeling and simulations (Goodreau, 2006), but the structure of sexual contact networks has been found to directly influence pathogen phylogenies (Robinson et al., 2013). Phylogenetic analysis of HIV type 1 sequences has revealed distinct grouping patterns based on risk behaviors (Holmes et al., 1995). The expectation of homophily is so strong, that its disruption, e.g. the presence of a self-reported heterosexual risk group individual in a cluster exclusively composed of MSMs has been used as a marker of undisclosed/incomplete risk factor reporting Ragonnet-Cronin et al. (2018a). Therefore, when subject-level attributes are available, homophily is an expected and desired feature of the network.

To assess the performance of an AUTO-TUNEd optimized threshold using degree-weighted homophily, we first evaluated a CRF07_BC network with national survey data from China Ge et al. (2021) . Each of the 8178 pol sequences was annotated with a transmission risk factor: heterosexual contact (Hetero), people who use injection drugs (PWID), or men who have sex with men (MSM), among other attributes.

When we analyze the dataset with AUTO-TUNE, local maxima of AUTO-TUNE scores were achieved with 0.0076 sub/site and 0.0019 sub/site thresholds, at scores 1.137 and 1.030, respectively. Notably, the DWH scores for PWID exhibited a significant surge at these thresholds, indicating a robust pattern of increased PWID homophily even when relatively low scoring. The close proximity of AUTO-TUNE scores and the consistent rise in PWID homophily at 0.0076 and 0.0019 thresholds suggest comparable performance at these thresholds compared to the default 0.015 threshold, suggesting that these thresholds might be more effective in representing homophilic relationships in this network. At each threshold—0.015, 0.0076, and 0.0019—all DWH scores for the risk groups (MSM, Hetero, and PWID) lie outside their respective panmictic ranges. This consistently indicates non-random mixing and attribute-based homophily across the network. Detailed results are in Table 7 and Table 8. The observation of significant homophily among PWID at lower thresholds identified by AUTO-TUNE (0.0076 and 0.0019 subs/site) is consistent with epidemiological characteristics of PWID outbreaks, such as the Drug-Resistant Subtype C Outbreak in Scotland, where rapid transmission led to low genetic diversity within clusters. (Ragonnet-Cronin et al., 2018b)

**Table 6.**
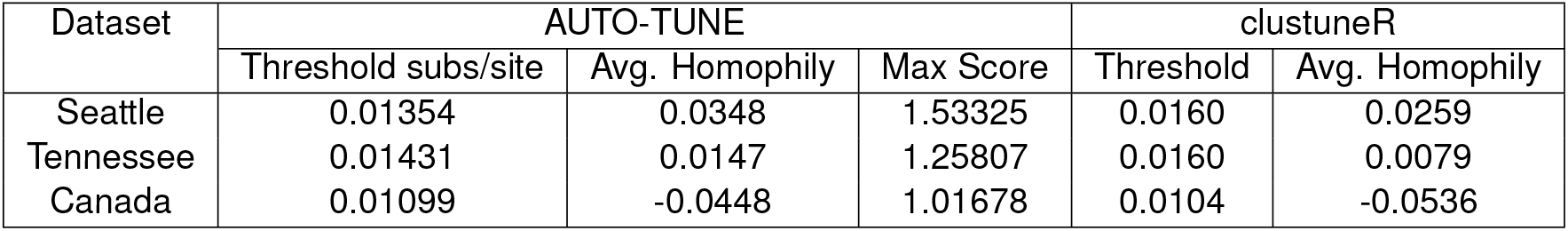
clustuneR Comparison.

**Table 7.** CRF07_BC Nodes Count at Different Thresholds.

| Threshold sub-s/site | AUTO-TUNE Score | Nodes | PWID | MSM | Hetero |
| --- | --- | --- | --- | --- | --- |
| 0.015 | 0.029 | 5923 | 559 | 3371 | 1993 |
| 0.0076 | 1.1369 | 3537 | 236 | 2271 | 1030 |
| 0.0019 | 1.0303 | 1654 | 151 | 1075 | 428 |

**Table 8.** Panmictic Ranges for CRF07_BC DWH at Different Thresholds.

| Threshold subs/site | Risk Group | DWH (Panmictic Range) |
| --- | --- | --- |
| 0.015 | MSM | 0.211(-0.213, -0.085) |
|  | Hetero | 0.133(-0.190, -0.087) |
|  | PWID | 0.168(-0.091, 0.002) |
| 0.0076 | MSM | 0.237(-0.240, -0.120) |
|  | Hetero | 0.185(-0.211, -0.100) |
|  | PWID | 0.401(-0.081, -0.005) |
| 0.0019 | MSM | 0.292(-0.280, -0.146) |
|  | Hetero | 0.250(-0.256, -0.093) |
|  | PWID | 0.445(-0.129, -0.012) |

### I. Comparison with clustuneR

We benchmarked AUTO-TUNE versus `clustuneR` Chato et al. (2020), which employs the recency of sample collection or diagnosis as individual-level weights in a predictive model to estimate the growth of HIV clusters. The thresholds deemed optimal by `clustuneR` were found by a grid-search for the minimum GAIC (generalized Akaike Information Criterion) across candidate distances between 0 and 0.04 in steps of 8 *×* 10^−4^. GAIC is the difference between a null model that is only influenced by cluster size, and a weighted model that includes individual-level attributes among known cases in the cluster. Using the minimum GAIC metric, it was found that 0.016(*±*0.5 *×* 10^−4^) was the optimal threshold for Tennessee and Seattle, and 0.0104 for Northern Alberta.

In contrast, AUTO-TUNE does not incorporate any attribute data in its scoring heuristic. Instead, it relies on clustering metrics constructed purely from pairwise distances between sequences. Using nearly same datasets analyzed by `clustuneR` (Chato et al., 2020), AUTO-TUNE found the thresholds with the highest scores to be 0.01872 for Middle Tennessee, 0.01538 for Seattle, and 0.01201 for Northern Alberta. Table 6. We use the adjective “nearly” because we were not able to exactly match the number of sequences analyzed in Chato et al. (2020) by obtaining the referenced GenBank accession number and our best-effort intepretation of the filtering steps.

Both methods agree that there is a qualitative relationship of Northern Alberta *<* Seattle ∼ Tennessee for distance thresholds. AUTO-TUNE thresholds, while not optimal in the GAIC sense all yield improvements over the null model, hence they are qualitatively similar to `clustuneR` (Figure 4 in Chato et al. (2020)). AUTO-TUNE is notably faster in computation than clustuneR due to the fact that AUTO-TUNE only clusters based on pairwise distances rather than inferring a maximum-likelihood phylogeny. For example, the entire pipeline for the Seattle dataset took less than `16` seconds on an Apple M1 Max. Alternatively, the tree inference step alone with `clustuneR` takes several hours to complete. Because the methods optimize very different objectives and `clustuneR` makes use of additional data, broad agreement between the inferred thresholds is encouraging.

### J. The Effect of Subsampling on Optimal Thresholds and AUTO-TUNE Scores

To address the challenges of applying network inference algorithms to incompletely sampled datasets, this study includes a focused evaluation of AUTO-TUNE’s performance across varying data densities. Given logistical limitations, obtaining a fully sampled HIV transmission network is often infeasible. Therefore, we label a dataset as ‘full’ to serve as a closest approximation of a fully sampled network. Using the selected dataset as a benchmark, we assess AUTO-TUNE’s adaptability and robustness when applied to sparser datasets, a prevalent issue in real-world settings.

Since the (Rhee et al., 2019) dataset exhibited a clear optimal peak, we used the dataset for analysis, and randomly sampled 10 times from the entire dataset at 25%, 50%, and 75% each. The original full dataset confidently determined 0.01699 (AUTO-TUNE score 1.9998).

Sampling at 25% yielded a mean top threshold of 0.021509, median at 0.019765, and standard deviation of 0.004388 1. 50% yielded 0.018581 and 0.01871 mean and median, respectively with a standard deviation of 0.001629. Finally, 75% calculated mean is approximately 0.017403, with a median of approximately 0.01699. The standard deviation was 0.000924.

As the dataset becomes sparser due to subsampling, the algorithm tends to select higher distance thresholds. This phenomenon can be understood by considering the effect of reduced sampling density on the network topology. Sparse datasets naturally result in less interconnected clusters. To capture a comparable level of network connectivity as in denser datasets, higher distance thresholds are necessary. This is evidenced by the observed mean thresholds: 0.021509 at 25%, 0.018581 at 50%, and 0.017403 at 75%. The standard deviations also narrow as the sampling density increases, corroborating the increased precision of the threshold selection in denser datasets.

As the proportion increased from 25% to 50% and 75%, observable shifts were also noted in the mean, median, and standard deviation of the AUTO-TUNE scores. At 25%, the mean and median scores were 1.5585 and 1.5014 respectively, with a standard deviation of 0.3568. At 50%, both mean and median scores significantly increased to 1.8171 and 1.9191 respectively, and the standard deviation dropped to 0.2482. Upon reaching an AUTO-TUNE of 75%, the mean and median scores rose further to 1.9870 and 1.9997 respectively, while the standard deviation shrank substantially to 0.0364, indicating higher consistency in scores.

Next to determine how well subsampled datasets aligned with the full dataset, we used two primary outcomes to gauge this concordance: the proportion of nodes that remained clustered after subsampling and the proportion of singletons from the original network that clustered in the subsampled networks.

We observed a consistent increase in the proportion of nodes that remained clustered from the 0.015 sub/site threshold to the AUTO-TUNE threshold for each respective subsampling proportion, with 25% subsampling being the most profound difference rising from a roughly 80%-86% interquartile range (IQR) for 0.015 threshold to a 90% 96% IQR for AUTO-TUNE, which indicates that the AUTO-TUNE thresholds retain a higher degree of stability in the network’s structure across sampling density (Please see Figure 2, Panel A).

Since the thresholds inferred by AUTO-TUNE for the sub-sampled networks were larger than the “fully” sampled network, we also measured the impact of thresholding on the network’s nodes that were originally singletons. Across all variations in subsampling rates, the proportion of sampled singletons that clustered all maintained low IQRs (See Figure 2, Panel B). This implies that while AUTO-TUNE is effective in maintaining the core structure of the network, it does not significantly alter the clustering of nodes that were singletons in the full dataset.

As the sample proportion increased, an upward trend was noted in average AUTO-TUNE scores. Additionally, the standard deviation reduced significantly when increasing sample proportion. This implies that as sampling becomes denser, AUTO-TUNE will become more confident in determining the optimal threshold for a particular dataset.

## Discussion

AUTO-TUNE addresses the challenge of selecting an appropriate genetic distance threshold to construct HIV transmission networks by implementing a heuristic scoring system. This system is predicated on two key features of networks generated by candidate genetic distance thresholds: a high number of clusters and the absence of a giant component. Few small clusters indicate an excessively low threshold, while a giant cluster comprising numerous sequences signals an overly high threshold. The efficacy of AUTO-TUNE is evidenced by its ability to select thresholds that yield higher quality clustering, as demonstrated by improved Degree-Weighted Homophily (DWH) scores across various datasets, epidemic contexts, and risk groups. Furthermore, AUTO-TUNE thresholds not only matched but often outperformed those manually selected in prior studies, thus underlining the benefits of a more systematic, automated, and data-responsive approach.

For example, the results of our study suggest that AUTO-TUNE, which relies solely on clustering metrics from pair-wise distances, could be an effective alternative to other distance-based methods, such as `clustuneR` while less time-consuming and possessing a gentle learning curve, which makes it easy to use by personnel not specialized in bioinformatics and computer science. Furthermore, the simplicity of the method without compromising results represents an advantage over phylogenetic methods where, in addition to the calculation of genetic distances, it must also determine a support/distance threshold where a rationale for the selection of these thresholds is rarely provided (Junqueira et al., 2019).

AUTO-TUNE generated thresholds for all three examined datasets (Middle Tennessee, Seattle, and Northern Alberta) that outperformed `clustuneR` using DWH on 3-year collection date windows across all three datasets. This indicates that even without incorporating attribute data, AUTO-TUNE’s scoring heuristic could provide reliable thresholds for HIV clusters. However, for the determination of the optimal genetic distance threshold, time-related and context-specific factors might need to be considered if there is no significant score for any one candidate threshold, especially if there are multiple peaks. For example, during HIV outbreaks in injection drug users (that usually occur over several months), it may be more appropriate to use the shorter genetic distance threshold (Peters et al., 2016; Campbell et al., 2017) between multiple high-scoring thresholds. On the contrary, larger and more extended epidemics over time exhibit a tendency toward larger genetic distance thresholds in order to capture transmission than younger epidemics and less densely sampled epidemic investigations (Patil et al., 2022; Leung et al., 2019; Di Giallonardo et al., 2021).

Our evaluation of publications citing HIV-TRACE revealed the largely qualitative determination of distance thresholds. This approach may result in less accurate or suboptimal thresholds due to a lack of systematic analysis. In contrast, AUTO-TUNE offers a more systematic and granular approach to threshold selection, with our findings demonstrating that even minor adjustments to the distance can drastically change the score. Therefore, using AUTO-TUNE could potentially improve the quality of HIV clustering and transmission network studies.

The Degree-Weighted Homophily (DWH) evaluation showed that AUTO-TUNE could improve network quality based on specific attributes, such as risk factor, which is an important part of HIV studies and informing prevention measures (Potterat et al., 2002; Fujimoto et al., 2021). For example, the use of AUTO-TUNE resulted in an increased DWH among the MSM, Hetero, and PWID groups when analyzing a CRF07_BC network. Additionally, the results from the Rhee et al. dataset also demonstrated AUTO-TUNE’s ability to improve DWH geographically, enhancing the network’s ability to accurately reflect transmission dynamics. However, in contexts with overlapping risk factors, the interpretation of these improvements requires caution. The complexities of risk group interactions mean that applying AUTO-TUNE’s thresholds should be tailored to the specific epidemiological setting to ensure accurate modeling of HIV transmission networks.

Our analysis of AUTO-TUNE’s performance on subsamples of a dataset revealed its sensitivity to sample size. The results indicated a correlation between increased sample size and higher average AUTO-TUNE scores, as well as lower score variability. This suggests that denser sampling could enhance AUTO-TUNE’s ability to determine the optimal threshold for a dataset. Further studies might be needed to establish the minimum sample size required for reliable threshold determination.

### K. When a Score is Below 1.9

In some cases, multiple scores at different thresholds could suggest the presence of inherently different scales in the network. For instance, if a network combines both global and local transmission patterns, AUTO-TUNE may produce more than one high score, reflecting these different scales. This was observed in a study on HIV-1 CRF07_BC transmission networks in China, where two distinct clusters, 07BC_N and 07BC_O, showed different transmission routes and geographic concentrations (Ding et al., 2022). Such network complexities could mean that different thresholds might offer more accurate insights into subpopulations or transmission dynamics. The use of AUTO-TUNE, while offering a method for automated threshold selection, may not always provide a single, decisive score that unambiguously determines the optimal threshold. In certain situations, such as datasets with lower sampling densities or those reflecting heterogenous dynamics within an epidemic, several candidate thresholds may yield similar AUTO-TUNE scores, making it difficult to single out one as the clear-cut ‘optimal’ threshold. In these scenarios, the process of threshold selection becomes more nuanced and requires a deeper analysis. The plot of AUTO-TUNE scores across candidate thresholds can serve as a valuable tool in these cases. For instance, researchers could identify a range of thresholds that all produce similar scores, suggesting that the specific choice of threshold within this range may not significantly impact the resulting network. Moreover, combining AUTO-TUNE with the DWH measure can enhance the interpretation of such plots. By considering how assortativity changes across the range of candidates, researchers can make more informed decisions about the appropriate choice. If there is a certain threshold at which the DWH measure noticeably changes for an attribute of interest, this could suggest a meaningful shift in the network structure that would be worth considering when selecting a threshold. The symbiotic approach of combining AUTO-TUNE scores, DWH measure, and visual analysis of score plots provides a more nuanced method for threshold selection when no clear optimal threshold emerges from the AUTO-TUNE scores alone.

The AUTO-TUNE methodology has several limitations. First, even though it provides the advantage of operating without the need for metadata, the size and the subgenomic region analyzed may affect the accuracy of transmission inference (Junqueira et al., 2019). Second, our analysis of AUTO-TUNE’s performance on subsamples of a dataset revealed its sensitivity to sample size, as the performance of the method can be affected by sampling density, improving the reliability of the test as the sampling density increases (figure X). However, our results were consistent with previous studies, which have suggested an optimal sampling density of 50 − 70% for HIV-1 cluster analysis (Novitsky et al., 2014). Third, even when it provides an insight of the optimal threshold to analyze a network, the supplied information might still need validation by experts, especially when no clear threshold is identified. In this case, it has been recommended to combine genetic data with clinical and sociodemographic information for a better characterization of the network structure. Finally, the performance of the method needs to be assessed in pathogens different from HIV, leading to opportunities for future research.

## Conclusion

AUTO-TUNE operates solely utilizing genetic sequence data to ascertain a decisive threshold. It employs a scoring heuristic, which is based on the number of clusters produced by a pairwise distance threshold and the ratio of the largest cluster to the second largest across a range of possible thresholds using sliding windows.

A key advantage of this approach is its autonomy from supplementary data. When a patient receives an HIV diagnosis, data collection protocols can greatly vary, and additional data are not always available or consistent. However, by leveraging only genetic sequence data, AUTO-TUNE eliminates the need for such information in some cases, and at minimum serves as a preliminary assessment of candidate thresholds.

Consequently, AUTO-TUNE’s performance is consistently controlled, irrespective of the fluctuations seen in data collection protocols after an HIV diagnosis. This level of adaptability demonstrates its suitability for integration into various contexts related to HIV, and possibly other viral cluster detection and response protocols. This versatility underscores the strong methodological foundation of AUTO-TUNE and its potential utility.

## Author Contributions

SW: Project administration, Conceptualization, Methodology, Writing – original draft; SLKP: Project administration, Conceptualization, Methodology, Writing – original draft, review & editing; DJ: Software; VDC: Data curation, Writing – original draft; HV: Visualization; SÁ-R: Writing – review & editing; AJLB: Writing – review & editing; JOW: Writing – review & editing.

## Supplementary Note 1: Figure captions

